## Supplementary material for "Ω-Loop mutations control dynamics of the active site by modulating the hydrogen-bonding network in PDC-3 β-lactamase": Figure 3 source data 1

**Table S1** MFPT (ns) of transitions between metastable states in wild-type PDC-3.

| state | 1 | 2 | 3 | 4 | 5 | 6 | 7 |
| --- | --- | --- | --- | --- | --- | --- | --- |
| 1 | 0.0 ± 0.0 | 38.6 ± 11.4 | 5089.2 ± 686.9 | 3428.7 ± 412.6 | 2586.9 ± 329.1 | 2232.9 ± 294.7 | 96.0 ± 15.6 |
| 2 | 1033.1 ± 234.0 | 0.0 ± 0.0 | 5079.7 ± 685.0 | 3415.4 ± 411.6 | 2571.2 ± 328.7 | 2217.1 ± 294.2 | 73.3 ± 14.0 |
| 3 | 3323.5 ± 411.5 | 2217.2 ± 262.6 | 0.0 ± 0.0 | 176.6 ± 48.4 | 577.1 ± 77.0 | 236.3 ± 53.9 | 1529.9 ± 181.7 |
| 4 | 3330.7 ± 399.6 | 2225.3 ± 243.9 | 1722.1 ± 337.1 | 0.0 ± 0.0 | 441.7 ± 51.3 | 104.4 ± 23.1 | 1503.7 ± 177.1 |
| 5 | 3309.7 ± 389.5 | 2205.6 ± 228.2 | 3060.5 ± 455.0 | 1251.2 ± 136.2 | 0.0 ± 0.0 | 23.5 ± 4.4 | 1448.8 ± 173.3 |
| 6 | 3315.0 ± 390.0 | 2210.8 ± 229.0 | 2998.6 ± 452.8 | 1188.6 ± 132.6 | 270.0 ± 26.1 | 0.0 ± 0.0 | 1455.7 ± 173.7 |
| 7 | 1898.7 ± 304.7 | 798.7 ± 88.0 | 5119.7 ± 662.7 | 3425.1 ± 398.5 | 2552.3 ± 324.3 | 2198.0 ± 289.6 | 0.0 ± 0.0 |

**Table S2** MFPT (ns) of transitions between metastable states in V211A variant.

| state | 1 | 2 | 3 | 4 | 5 | 6 | 7 |
| --- | --- | --- | --- | --- | --- | --- | --- |
| 1 | 0.0 ± 0.0 | 941.5 ± 67.8 | 722.7 ± 51.0 | 1913.7 ± 162.7 | 158.9 ± 8.1 | 1931.0 ± 178.5 | 34.2 ± 4.3 |
| 2 | 2281.6 ± 239.2 | 0.0 ± 0.0 | 133.3 ± 12.8 | 1206.4 ± 139.8 | 155.9 ± 17.1 | 1363.8 ± 154.6 | 489.8 ± 49.4 |
| 3 | 2362.1 ± 241.8 | 408.9 ± 37.0 | 0.0 ± 0.0 | 958.4 ± 117.7 | 238.4 ± 27.2 | 1091.0 ± 139.7 | 570.6 ± 58.6 |
| 4 | 2542.9 ± 244.8 | 511.4 ± 46.1 | 104.3 ± 7.8 | 0.0 ± 0.0 | 424.3 ± 36.3 | 767.0 ± 102.0 | 752.1 ± 65.7 |
| 5 | 1981.4 ± 228.5 | 671.4 ± 61.3 | 452.7 ± 44.0 | 1650.5 ± 159.9 | 0.0 ± 0.0 | 1684.3 ± 174.0 | 191.5 ± 22.2 |
| 6 | 2665.8 ± 253.9 | 764.7 ± 52.8 | 304.7 ± 23.5 | 822.0 ± 86.0 | 561.0 ± 48.1 | 0.0 ± 0.0 | 876.8 ± 78.0 |
| 7 | 1704.1 ± 212.4 | 883.1 ± 66.9 | 664.5 ± 49.9 | 1856.1 ± 161.8 | 100.9 ± 4.5 | 1874.8 ± 177.6 | 0.0 ± 0.0 |

**Table S3** MFPT (ns) of transitions between metastable states in V211G variant.

| **state** | **1** | **2** | **3** | **4** | **5** | **6** | **7** | **8** |
| --- | --- | --- | --- | --- | --- | --- | --- | --- |
| 1 | 0.0 ± 0.0 | 549.3 ± 32.0 | 2081.8 ± 266.3 | 799.0 ± 66.5 | 176.5 ± 9.7 | 147.0 ± 7.5 | 15.1 ± 3.1 | 355.0 ± 19.2 |
| 2 | 3010.2 ± 486.4 | 0.0 ± 0.0 | 2180.4 ± 257.5 | 835.0 ± 60.7 | 65.0 ± 5.7 | 134.6 ± 6.1 | 405.0 ± 29.0 | 231.3 ± 16.7 |
| 3 | 3032.2 ± 484.5 | 668.3 ± 36.7 | 0.0 ± 0.0 | 98.6 ± 14.1 | 374.5 ± 19.7 | 84.4 ± 7.1 | 456.4 ± 32.9 | 248.2 ± 15.9 |
| 4 | 3055.3 ± 485.8 | 638.4 ± 35.6 | 1320.0 ± 215.7 | 0.0 ± 0.0 | 353.1 ± 18.4 | 48.3 ± 4.1 | 461.8 ± 32.2 | 188.6 ± 13.9 |
| 5 | 2894.2 ± 484.4 | 300.8 ± 25.4 | 2139.0 ± 257.2 | 804.0 ± 60.7 | 0.0 ± 0.0 | 105.5 ± 5.2 | 291.9 ± 25.5 | 250.5 ± 16.4 |
| 6 | 3019.9 ± 485.6 | 556.7 ± 34.3 | 1972.4 ± 255.0 | 613.7 ± 56.6 | 272.5 ± 16.7 | 0.0 ± 0.0 | 411.2 ± 30.8 | 155.5 ± 11.3 |
| 7 | 2435.1 ± 464.7 | 515.4 ± 32.0 | 2076.5 ± 258.4 | 778.1 ± 62.7 | 142.1 ± 8.6 | 114.7 ± 6.5 | 0.0 ± 0.0 | 323.5 ± 18.7 |
| 8 | 3217.9 ± 486.7 | 631.2 ± 36.3 | 2165.4 ± 256.7 | 782.3 ± 59.9 | 399.8 ± 19.7 | 143.2 ± 5.5 | 611.2 ± 33.7 | 0.0 ± 0.0 |

**Table S4** MFPT (ns) of transitions between metastable states in G214A variant.

| **state** | **1** | **2** | **3** | **4** | **5** | **6** | **7** | **8** |
| --- | --- | --- | --- | --- | --- | --- | --- | --- |
| 1 | 0.0 ± 0.0 | 51.2 ± 14.8 | 5343.0 ±1796.1 | 2598.2 ± 400.4 | 2063.5 ± 390.7 | 1152.6 ± 301.7 | 1591.0 ± 368.0 | 1667.2 ± 372.9 |
| 2 | 4009.0 ± 1661.6 | 0.0 ± 0.0 | 5018.9 ± 1739.6 | 2490.3 ± 400.6 | 1957.2 ± 389.7 | 1043.4 ± 301.2 | 1483.1 ± 367.8 | 1559.4 ± 372.8 |
| 3 | 6682.9 ± 2324.3 | 652.1 ± 233.4 | 0.0 ± 0.0 | 2615.3 ± 400.7 | 2089.0 ± 389.3 | 1162.8 ± 303.7 | 1608.2 ± 368.0 | 1685.1 ± 373.3 |
| 4 | 15106.5 ± 3589.7 | 9381.3 ± 2015.7 | 13741.1 ±3171.9 | 0.0 ± 0.0 | 1459.8 ± 122.9 | 687.2 ± 49.2 | 32.6 ±2.7 | 219.5 ± 12.2 |
| 5 | 14532.2 ± 3582.3 | 8810.9 ± 2004.4 | 13175.0 ±3162.7 | 1414.4 ±114.2 | 0.0 ± 0.0 | 108.1 ± 9.6 | 414.0 ± 43.1 | 436.5 ± 50.2 |
| 6 | 14270.1 ± 3580.2 | 8545.2 ± 1998.5 | 12905.0 ±3156.6 | 1352.6 ±120.4 | 660.6 ± 88.2 | 0.0 ± 0.0 | 346.4 ± 49.6 | 409.6 ± 57.2 |
| 7 | 15013.6 ± 3590.3 | 9288.4 ± 2015.8 | 13648.1± 3172.1 | 869.6 ± 83.1 | 1370.7 ± 122.7 | 595.4 ± 48.1 | 0.0 ± 0.0 | 145.9 ± 10.8 |
| 8 | 15021.8 ± 3589.9 | 9296.8 ± 2014.8 | 13657.1±3171.0 | 1065.8 ± 89.2 | 1327.9 ± 122.5 | 597.0 ± 46.5 | 95.1 ± 5.1 | 0.0 ± 0.0 |

**Table S5** MFPT (ns) of transitions between metastable states in G214R variant.

| **state** | **1** | **2** | **3** | **4** | **5** | **6** | **7** |
| --- | --- | --- | --- | --- | --- | --- | --- |
| 1 | 0.0 ± 0.0 | 141.0 ± 38.8 | 15986.1 ± 4905.3 | 24549.1 ± 7191.9 | 146.4 ± 25.5 | 359.3 ± 32.5 | 389.1 ± 34.0 |
| 2 | 11051.2 ± 2573.8 | 0.0 ± 0.0 | 15890.0 ± 4903.5 | 24453.0 ± 7192.4 | 59.1 ± 6.4 | 268.8 ± 18.5 | 291.3 ± 22.1 |
| 3 | 14842.1 ± 2801.5 | 3661.6 ± 742.9 | 0.0 ± 0.0 | 3726.9 ± 1518.2 | 2494.5 ± 678.4 | 1868.5 ± 645.9 | 1461.0 ± 539.6 |
| 4 | 15082.9 ± 2798.1 | 3902.3 ± 750.4 | 184.3 ± 40.0 | 0.0 ± 0.0 | 2735.2 ± 687.1 | 2109.3 ± 654.8 | 1701.6 ± 551.9 |
| 5 | 12107.2 ± 2638.7 | 949.7 ± 139.7 | 15810.8 ± 4904.1 | 24373.8 ± 7193.0 | 0.0 ± 0.0 | 189.3 ± 14.2 | 212.2 ± 18.5 |
| 6 | 12967.4 ± 2656.5 | 1793.2 ± 205.7 | 15626.3 ± 4900.6 | 24189.3 ± 7188.2 | 630.0 ± 60.6 | 0.0 ± 0.0 | 74.6 ± 5.0 |
| 7 | 13104.8 ± 2657.4 | 1924.4 ± 213.3 | 15478.5 ± 4898.7 | 24040.6 ± 7186.8 | 757.4 ± 69.7 | 128.4 ± 11.2 | 0.0 ± 0.0 |

**Table S6** MFPT (ns) of transitions between metastable states in E219A variant.

| **state** | **1** | **2** | **3** | **4** | **5** | **6** | **7** |
| --- | --- | --- | --- | --- | --- | --- | --- |
| 1 | 0.0 ± 0.0 | 1264.4 ± 687.4 | 2164.5 ± 410.6 | 1090.0 ± 295.7 | 1120.1 ± 351.4 | 87.1 ± 27.1 | 1144.0 ± 403.8 |
| 2 | 6228.8 ± 3113.1 | 0.0 ± 0.0 | 1901.1 ± 451.9 | 1108.5 ± 278.2 | 1092.4 ± 342.7 | 221.3 ± 43.6 | 1213.9 ± 359.7 |
| 3 | 12248.7 ± 4167.4 | 5437.4 ± 1621.2 | 0.0 ± 0.0 | 1020.7 ± 120.0 | 343.5 ± 81.3 | 278.9 ± 40.1 | 780.5 ± 119.3 |
| 4 | 12745.9 ± 4185.6 | 6535.0 ± 1631.1 | 2243.0 ± 288.1 | 0.0 ± 0.0 | 465.2 ± 46.7 | 116.7 ± 10.0 | 467.7 ± 49.8 |
| 5 | 13096.8 ± 4198.9 | 6777.4 ± 1649.2 | 2053.5 ± 278.3 | 942.1 ± 76.4 | 0.0 ± 0.0 | 86.0 ± 6.6 | 507.0 ± 42.4 |
| 6 | 12353.9 ± 4325.9 | 6318.3 ± 1690.8 | 2265.3 ± 282.7 | 864.4 ± 94.0 | 378.7 ± 102.5 | 0.0 ± 0.0 | 394.1 ± 108.6 |
| 7 | 13069.7 ± 4204.1 | 6876.9 ± 1645.0 | 2408.4 ± 281.1 | 898.1 ± 68.6 | 462.6 ± 31.8 | 68.3 ± 2.7 | 0.0 ± 0.0 |

**Table S7** MFPT (ns) of transitions between metastable states in E219G variant.

| **state** | **1** | **2** | **3** | **4** | **5** | **6** | **7** |
| --- | --- | --- | --- | --- | --- | --- | --- |
| 1 | 0.0 ± 0.0 | 1212.6 ± 252.9 | 3392.5 ± 403.3 | 3616.1 ± 448.4 | 56.4 ±13.0 | 1900.0 ± 358.7 | 2209.6 ± 373.9 |
| 2 | 19628.6 ± 3525.0 | 0.0 ± 0.0 | 1841.2 ± 150.1 | 2069.3 ± 252.4 | 10472.6 ± 2517.6 | 351.1 ± 70.4 | 658.1 ± 94.9 |
| 3 | 23439.6 ± 3756.0 | 2847.1 ± 334.7 | 0.0 ± 0.0 | 2300.0 ± 239.1 | 14284.2 ± 2819.3 | 411.3 ± 21.5 | 40.2 ± 3.1 |
| 4 | 22897.1 ± 3752.1 | 2331.5 ± 323.7 | 1533.7 ± 111.1 | 0.0 ± 0.0 | 13741.8 ± 2816.4 | 53.8 ± 5.8 | 360.8 ± 28.4 |
| 5 | 5362.9 ± 1288.4 | 1120.3 ± 252.8 | 3300.0 ± 403.6 | 3523.6 ± 449.8 | 0.0 ± 0.0 | 1807.6 ± 359.8 | 2117.1 ± 374.8 |
| 6 | 22856.3 ± 3754.4 | 2284.0 ± 325.0 | 1393.1 ± 108.4 | 1557.8 ± 223.1 | 13701.0 ± 2818.0 | 0.0 ± 0.0 | 220.9 ± 23.5 |
| 7 | 23358.6 ± 3756.0 | 2765.7 ± 334.2 | 1022.1 ± 90.1 | 2221.6 ± 238.4 | 14203.3 ± 2819.4 | 333.1 ± 19.7 | 0.0 ± 0.0 |

**Table S8** MFPT (ns) of transitions between metastable states in E219K variant.

| **state** | **1** | **2** | **3** | **4** | **5** | **6** | **7** |
| --- | --- | --- | --- | --- | --- | --- | --- |
| 1 | 0.0 ± 0.0 | 10327.3 ± 2768.6 | 8262.0 ± 2573.0 | 7806.9 ± 2528.0 | 6498.0 ± 2229.7 | 1621.4 ± 415.5 | 1985.3 ± 487.8 |
| 2 | 3469.4 ± 601.1 | 0.0 ± 0.0 | 29.5 ± 3.2 | 371.5 ± 38.4 | 1746.2 ± 314.4 | 5371.0 ± 1051.2 | 5648.8 ± 1101.7 |
| 3 | 3415.8 ± 601.4 | 933.8 ± 130.4 | 0.0 ± 0.0 | 378.2 ± 40.9 | 1674.5 ± 310.5 | 5322.9 ± 1051.0 | 5601.1 ± 1101.4 |
| 4 | 3131.3 ± 593.1 | 2048.6 ± 274.6 | 574.5 ± 119.4 | 0.0 ± 0.0 | 1548.9 ± 352.7 | 4993.4 ± 1048.7 | 5268.7 ± 1100.8 |
| 5 | 2193.5 ± 502.0 | 3936.1 ± 672.5 | 1692.9 ± 417.4 | 1661.1 ± 457.9 | 0.0 ± 0.0 | 4360.4 ± 972.0 | 4654.3 ± 1018.5 |
| 6 | 3408.7 ± 724.0 | 14634.3 ± 3506.3 | 12628.5 ± 3323.2 | 12006.3 ± 3287.6 | 11162.4 ± 3016.3 | 0.0 ± 0.0 | 120.2 ± 15.3 |
| 7 | 3599.4 ± 725.8 | 14773.1 ± 3510.7 | 12769.0 ± 3327.0 | 12141.5 ± 3294.3 | 11311.0 ± 3017.2 | 219.9 ± 14.7 | 0.0 ± 0.0 |

**Table S9** MFPT (ns) of transitions between metastable states in Y221A variant.

| **state** | **1** | **2** | **3** | **4** | **5** | **6** | **7** |
| --- | --- | --- | --- | --- | --- | --- | --- |
| 1 | 0.0 ± 0.0 | 323.9 ± 141.3 | 2324.2 ± 436.6 | 820.0 ± 340.2 | 1603.8 ± 387.3 | 936.5 ± 358.4 | 804.9 ± 345.2 |
| 2 | 16651.3 ± 6505.9 | 0.0 ± 0.0 | 1753.5 ± 168.9 | 267.6 ± 46.2 | 1027.7 ± 103.1 | 319.6 ± 59.6 | 139.5 ± 44.4 |
| 3 | 18741.5 ± 6567.0 | 1389.5 ± 81.2 | 0.0 ± 0.0 | 374.8 ± 26.5 | 39.3 ± 4.1 | 401.3 ± 27.5 | 780.2 ± 46.8 |
| 4 | 18148.3 ± 6555.8 | 840.2 ± 69.2 | 1349.4 ± 134.1 | 0.0 ± 0.0 | 647.0 ± 63.4 | 193.5 ± 17.9 | 312.7 ± 25.5 |
| 5 | 18685.7 ± 6566.4 | 1329.9 ± 80.7 | 491.9 ± 61.7 | 336.5 ± 23.7 | 0.0 ± 0.0 | 322.2 ± 27.7 | 715.9 ± 46.5 |
| 6 | 18349.2 ± 6556.1 | 945.4 ± 66.4 | 1443.0 ± 127.8 | 257.4 ± 14.8 | 694.9 ± 63.0 | 0.0 ± 0.0 | 254.8 ± 19.2 |
| 7 | 18167.2 ± 6551.0 | 708.8 ± 55.2 | 1737.6 ± 139.7 | 332.8 ± 18.0 | 1004.2 ± 73.7 | 216.5 ± 15.6 | 0.0 ± 0.0 |

**Table S10** MFPT (ns) of transitions between metastable states in Y221H variant.

| **state** | **1** | **2** | **3** | **4** | **5** | **6** | **7** |
| --- | --- | --- | --- | --- | --- | --- | --- |
| 1 | 0.0 ± 0.0 | 1730.6 ± 255.7 | 826.5 ± 54.7 | 679.3 ± 93.9 | 383.8 ± 26.2 | 87.1 ± 10.8 | 425.7 ± 28.5 |
| 2 | 1222.7 ± 170.4 | 0.0 ± 0.0 | 726.1 ± 57.7 | 840.6 ± 105.7 | 578.4 ± 31.2 | 242.7 ± 15.5 | 329.6 ± 31.5 |
| 3 | 1835.4 ± 187.0 | 2304.3 ± 273.8 | 0.0 ± 0.0 | 1328.5 ± 113.8 | 611.6 ± 34.0 | 214.0 ± 12.8 | 30.4 ± 3.0 |
| 4 | 1245.7 ± 180.0 | 1962.2 ± 272.4 | 898.0 ± 54.5 | 0.0 ± 0.0 | 382.2 ± 26.0 | 118.4 ± 10.7 | 494.3 ± 29.0 |
| 5 | 1632.9 ± 187.2 | 2401.6 ± 275.3 | 856.7 ± 52.8 | 1057.4 ± 108.9 | 0.0 ± 0.0 | 18.2 ± 0.8 | 449.4 ± 27.3 |
| 6 | 1629.4 ± 187.2 | 2382.2 ± 274.9 | 811.5 ± 52.8 | 1072.4 ± 109.3 | 275.2 ± 19.2 | 0.0 ± 0.0 | 404.3 ± 27.3 |
| 7 | 1843.2 ± 187.2 | 2323.0 ± 274.0 | 421.3 ± 32.5 | 1331.9 ± 113.8 | 609.5 ± 33.6 | 212.7 ± 12.0 | 0.0 ± 0.0 |
